# The importance of residue-level filtering, and the Top2018 best-parts dataset of high-quality protein residues

**DOI:** 10.1101/2021.10.05.463241

**Authors:** Christopher J. Williams, David C. Richardson, Jane S. Richardson

## Abstract

We have curated a high-quality, “best parts” reference dataset of about 3 million protein residues in about 15,000 PDB-format coordinate files, each containing only residues with good electron density support for a physically acceptable model conformation. The resulting pre-filtered data typically contains the entire core of each chain, in quite long continuous fragments. Each reference file is a single protein chain, and the total set of files were selected for low redundancy, high resolution, good MolProbity score, and other chain-level criteria. Then each residue was critically tested for adequate local map quality to firmly support its conformation, which must also be free of serious clashes or covalent-geometry outliers. The resulting Top2018 pre-filtered datasets have been released on the Zenodo online web service and is freely available for all uses under a Creative Commons license. Currently, one dataset is residue-filtered on mainchain plus Cβ atoms, and a second dataset is full-residue filtered; each is available at 4 different sequence-identity levels. Here, we illustrate both statistics and examples that show the beneficial consequences of residue-level filtering. That process is necessary because even the best of structures contain a few highly disordered local regions with poor density and low-confidence conformations that should not be included in reference data. Therefore the open distribution of these very large, pre-filtered reference datasets constitutes a notable advance for structural bioinformatics and the fields that depend upon it.

The Top2018 dataset provides the first representative sample of 3D protein structure for which excellence of experimental data constrains the detailed local conformation to be correct for essentially all 3 million residues included. Earlier generations of residue-filtered datasets were central in developing MolProbity validation used worldwide, and now Zenodo has enabled anyone to use out latest version as a sound basis for structural bioinformatics, protein design, prediction, improving biomedically important structures, or other applications.

## Introduction

Our laboratory has emphasized the importance of residue-level as well as chain-level quality filtering of reference datasets as a foundation for model validation and for further bioinformatic structural studies. We began such work in the late 1990s when we introduced our flagship validation of all-atom contact analysis based on the Top100 dataset of reference protein chains, which in our own use we filtered at the residue level on any atomic B-factor >40^1^. We made available the list for those 100 chains and for all our subsequent, increasingly large reference datasets (8000 chains by 2013), but had to leave the application of B cutoffs to the user. After deposition of structure factors became required, our validations used explicit electron-density filters for map value and correlation coefficient at each atom, as well as B-factor, all-atom clash, and covalent-geometry filters^2^, but we still found no feasible mechanism for distributing all the coordinate files with residue-filter annotations.

Our current residue-level quality filtering process relies on extensive infrastructure, especially our developer team’s integration into the Phenix software project^3^. We also now manage the filtering information with a Neo4j graphical database^4,5^. We have switched to using a graphical database to store our reference data because sequence connectivity is modeled natively there (but cumbersome in relational databases), as are the cyclic graphs that define local structural motifs.

The recent breakthrough in our ability to distribute coordinate files in a residue-filtered mode has been enabled by two things. First is our realization that making residue-level quality filtering easily available is worth giving up user flexibility in setting filter thresholds. Second, and most important, is the Zenodo online service that hosts open access to very large, DOI-identified datasets^6^. We have now taken advantage of that venue to distribute our current best-parts datasets. This development allows other researchers to make full and proper use of our curated reference data without needing the expertise, infrastructure, and effort required to perform residue-level quality-filtering themselves.

Here we outline the production of this high-quality Top2018 (∼15,000-chain) protein dataset and announce the availability of two residue-level pre-filtered versions suitable for general use with little or no further modification. One set is residue-filtered on mainchain criteria and the other on both mainchain and sidechain criteria. Each set is available at 30%, 50%, 70%, and 90% sequence-identity levels. The filtered-out residues leave gaps in the chain, but the remaining high-reliability fragments are surprisingly long – mostly 20-30 residues or more.

### Chain selection

We assembled a set of high-quality, low-redundancy protein chains.

Chains are selected for consideration from the Protein Data Bank on the following criteria:

- Chain is protein
- Sequence length ≥ 38 residues
- Parent structure solved with x-ray crystallography
- Parent structure solved at better than 2.0Å resolution
- Parent structure has deposited structure factors
- Parent structure deposited on or before December 31, 2018

These chains are analyzed with our validation statistics, and chains that fail the following criteria are removed from consideration:

- MolProbity score < 2.0^7^
- < 3% of residues have Cβ deviations^8^
- < 2% of residues have covalent bond length outliers > 4 σ
- < 2% of residues have covalent bond angle outliers > 4 σ

The remaining chains are treated within their PDB-defined sequence-identity clusters, which are calculated weekly with MMseqs2^9^. From each cluster, we select the chain with the best (lowest) average of resolution and MolProbity score as the best-quality representative of that cluster.

The PDB provides homology clustering at several different levels of stringency. We prepared sets of chains at the 90, 70, 50, and 30% sequence-identity levels. 90% is the most permissive, allowing as much as 90% sequence homology between the representatives from different clusters. 30% is the most restrictive, grouping chains into fewer clusters (and thus fewer one-per-cluster representatives), with greater differences between clusters. We recommend 70% for general use as a good balance point between coverage and non-redundancy.

To enable all-atom contact analysis^1^ and produce the best quality chain possible, the parent structures have hydrogens added and N/Q/H flips performed by Reduce^10^. This is done prior to splitting the structure into chains, so that hydrogen bonding networks and other contacts are kept intact within the asymmetric unit.

### Residue-level filtering

While the selected chains are of good overall quality, this does not guarantee that all residues in them are modeled at high quality with high confidence. Therefore, we apply a residue-level filtering process. Figure 1A shows 5Lp0^11^ typical Top2018 chain. The mainchain core is well resolved and broken only by 3 individual clashes. The few serious errors are concentrated in just two short surface regions of poor density.

**Figure 1:**
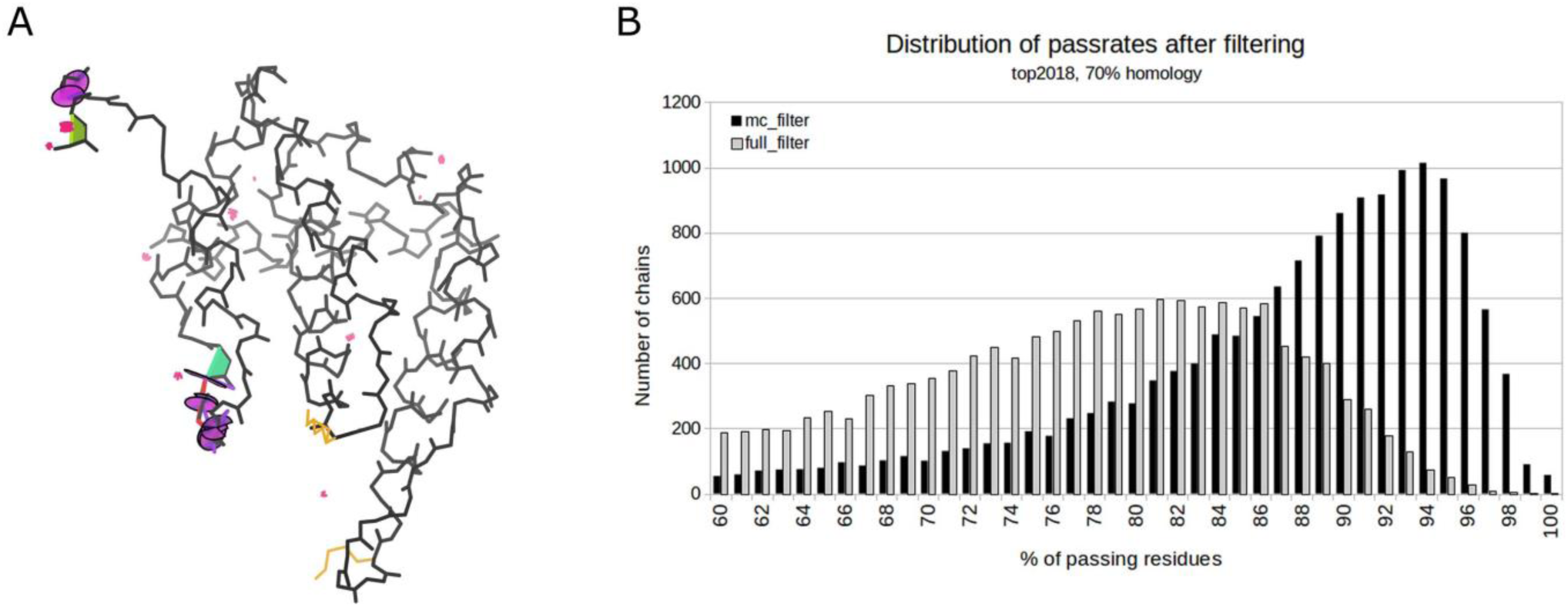
Distributions of model quality in high-resolution structure. (A) 5Lp0 demonstrates a typical distribution of structure quality for models included in the Top2018. Most of the model is reliable and free from outliers, but two short regions contain a concentration of significant errors, including the clashes (clusters of red spikes) and covalent-geometry outliers that trigger omissions, and the conformation outliers that also occur in these unreliable regions such as *cis*-peptides (green trapezoids) and CaBLAM outliers (magenta lines and purple wheels). (B) Distribution of % passing residues in Top2018 after mainchain (solid bars) and full-residue filtering (open bars).

Two different residue-filtered sets were created, one filtered just on the mainchain and one filtered on the full residue, including the sidechain. Mainchain filtering considers the atoms N, Cα, C, O, and Cβ. Cβ is included with the mainchain atoms since its ideal position is determined solely from other mainchain atom positions. Full-residue filtering considers all mainchain and sidechain heavy atoms. Hydrogen atoms are used for all-atom contact analysis, but not for fit-to-map analyses, as their signal in the map is weak or absent.

For a residue to be included in the final dataset, all atoms under consideration must meet the following criteria:

- B-factor < 40
- Real-space correlation coefficient (RSCC) > 0.7
- 2mFo-DFc map value at atom position > 1.2 σ (This level is represented by gray contours in Figures 2, 4, 5 and 6.)
- No covalent geometry outliers > 4 σ involving those atoms
- No steric clashes (overlaps ≥ 0.4 Å) involving those atoms
- No alternate conformations for those atoms

**Figure 2:**
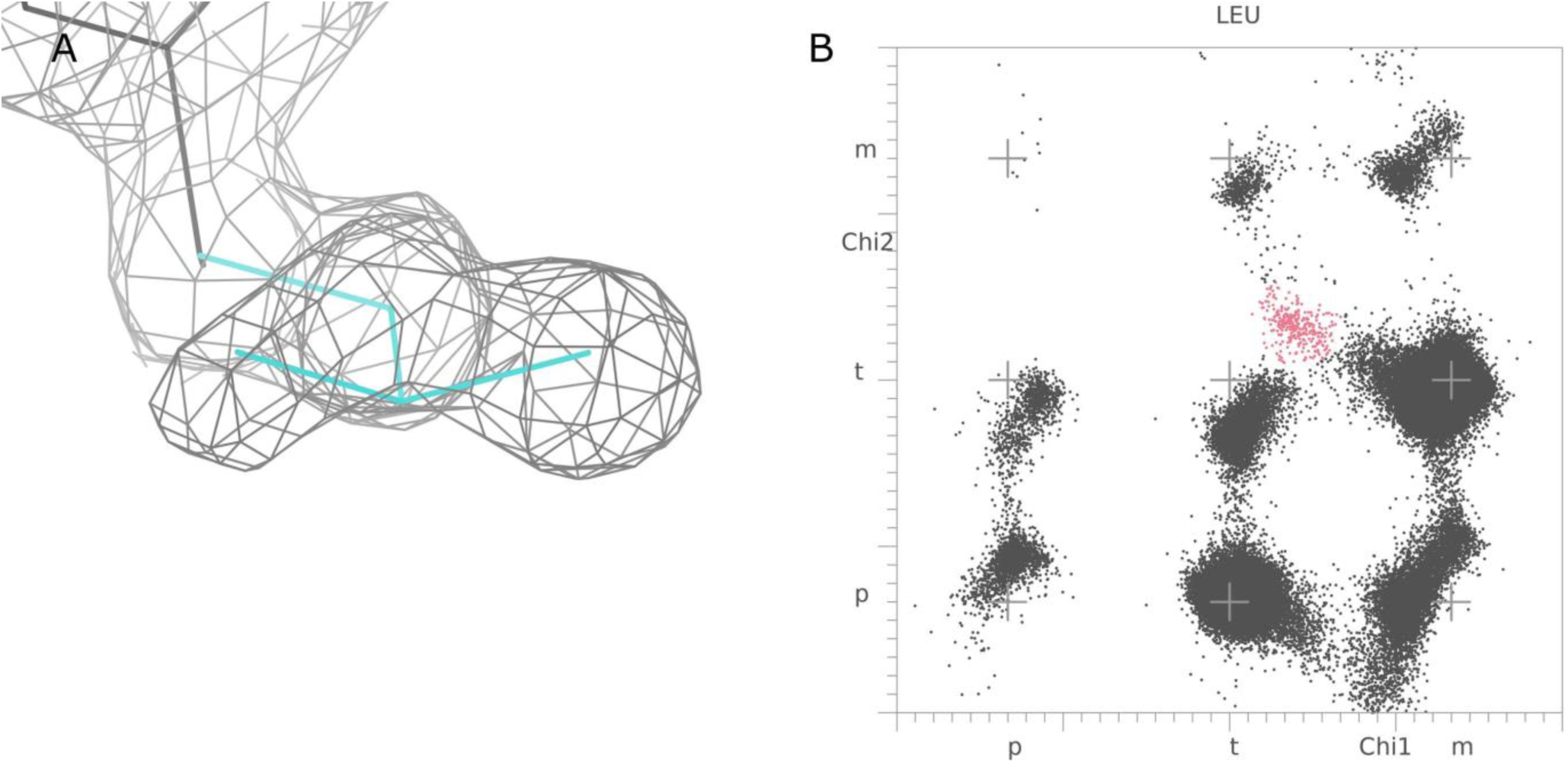
A systematic-error Leucine conformation. (A) Leucine has a common “imposter” conformation rotated 30-40° in χ1 and 140-150° in χ2 from the very common **tp** conformation, which keeps Cδ atoms in about the same peaks. Despite the sidechain geometry not fitting the shape of the density, the atoms are often within the gray 1.2σ contour well enough to pass our filters. (B) Even after filtering, a cluster of this outlier is clearly visible in pink to the upper right of the **tt** rotamer. The datapoints near χ2 zero in the **mp** bin are a similar systematic error fit backwards from the very common **mt** rotamer.

**Figure 3:**
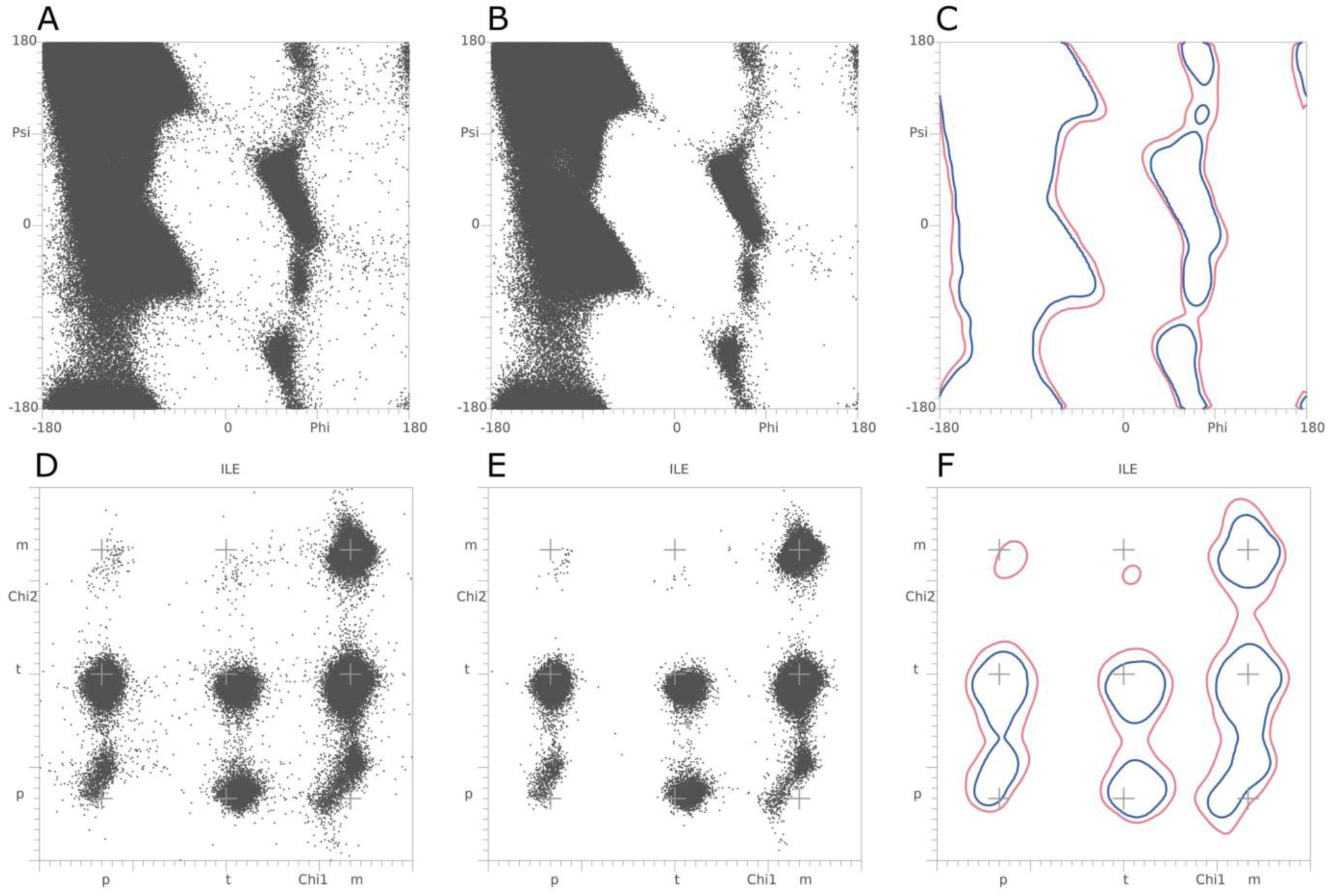
Ramachandran and rotamer distributions before and after residue filtering. (A) Ramachandran distribution for general-case residues in Top2018 70% sequence identity set before residue-level filtering. (B) Mainchain filtering cleans up distribution edges and removes many outliers. (C) 0.2% allowed/outlier contour boundary before (red) and after (blue) filtering. (D) χ1/χ2 rotamer distribution for isoleucine before filtering. (E) Full-residue filtering cleans up distribution edges and prevents very rare, highly strained conformations from being defined as rotamers. (F) 0.3% allowed/outlier contour boundary before (red) and after (blue) filtering.

**Figure 4:**
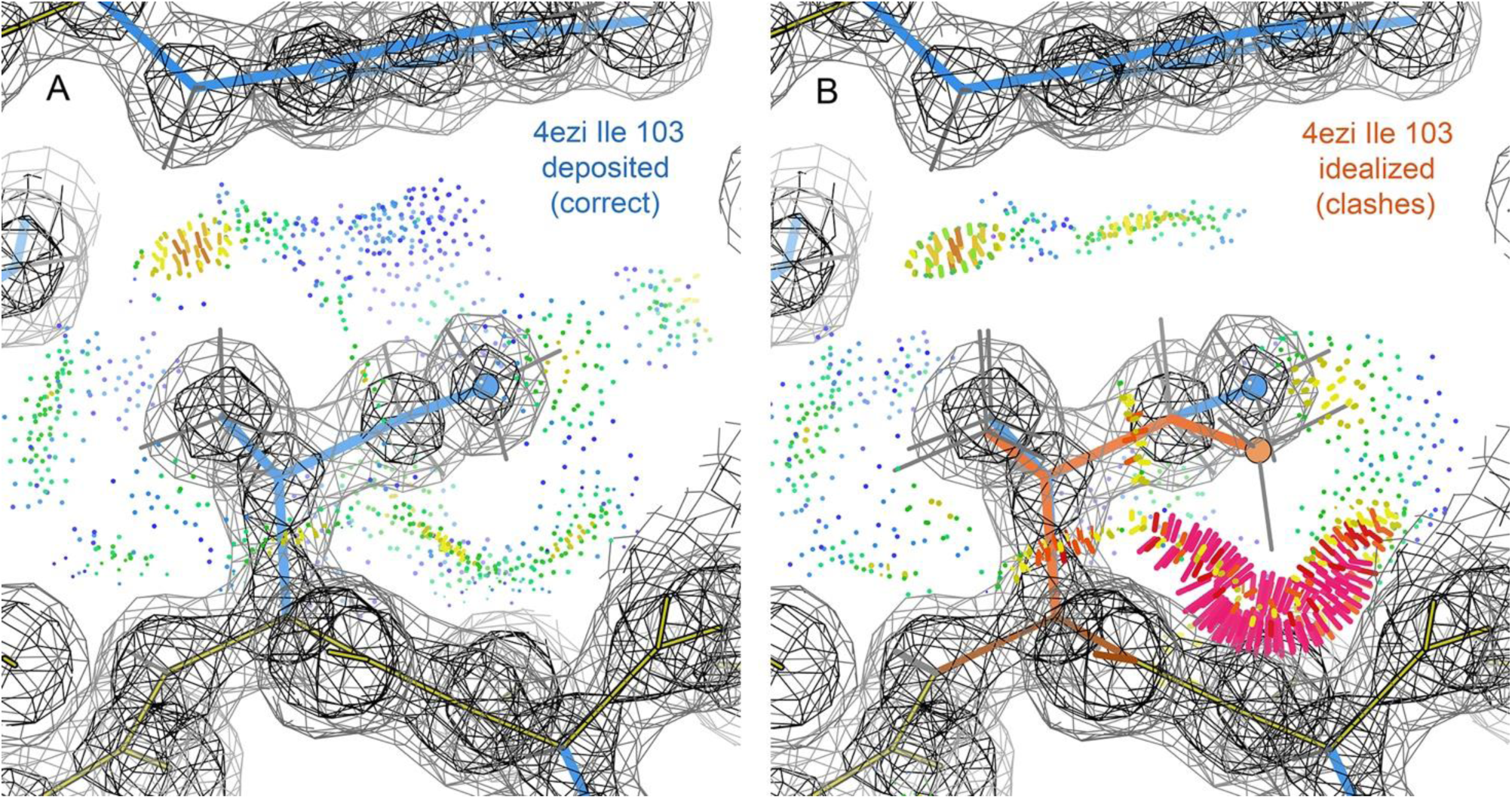
The rationale for a genuine but highly strained rare conformation. (A) The unambiguously correct **tm** model for 4ezi Ile 103 at 1.15Å, with a separate, strong density peak at every atom and good van der Waals contacts all around the sidechain (green and blue dots) and one small overlap (yellow and orange). The Ile Cδ atom is highlighted with a blue ball. (B) For comparison, the “ideal” **tm** conformation at 180° χ1, -60° χ2 is overlaid (in orange, with orange ball on Cδ), which has an extremely large clash (hotpink spikes). Gray contour represents 1.2σ map level used by filters.

**Figure 5:**
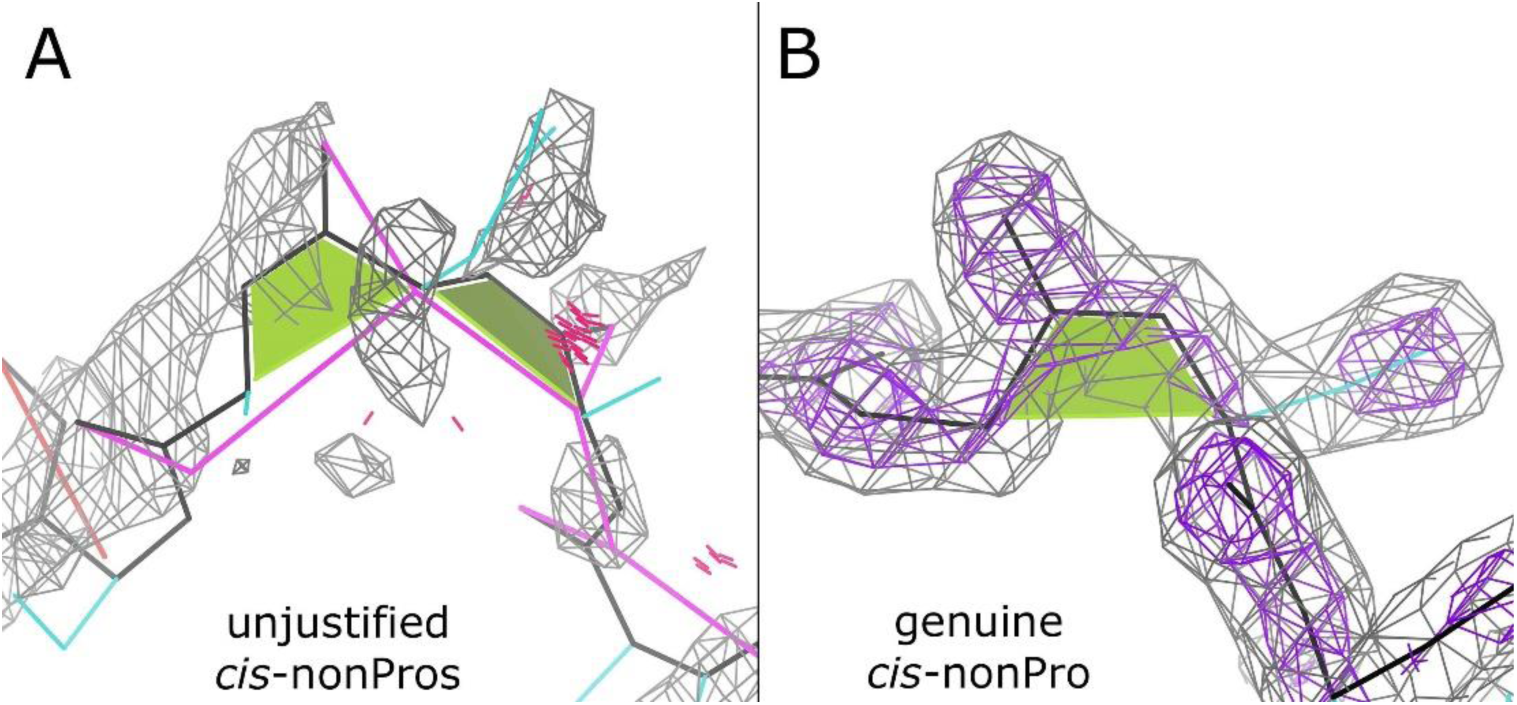
Non-proline *cis*-peptides (flagged by green trapezoids (A) A double *cis*-nonPro modeled into a loop region of poor density in 4rm4 residues 170-172. Real *cis*-nonPro never occur sequentially, and this is not a reasonable interpretation of the map. These residues are removed in filtering. (B) A genuine *cis*-serine, residue 275 of 6btf. It passes our quality criteria, and is included in the structure after filtering. The 1.6Å density is persuasive and well fit by the model. Gray contour represents 1.2σ map level used by filters.

**Figure 6:**
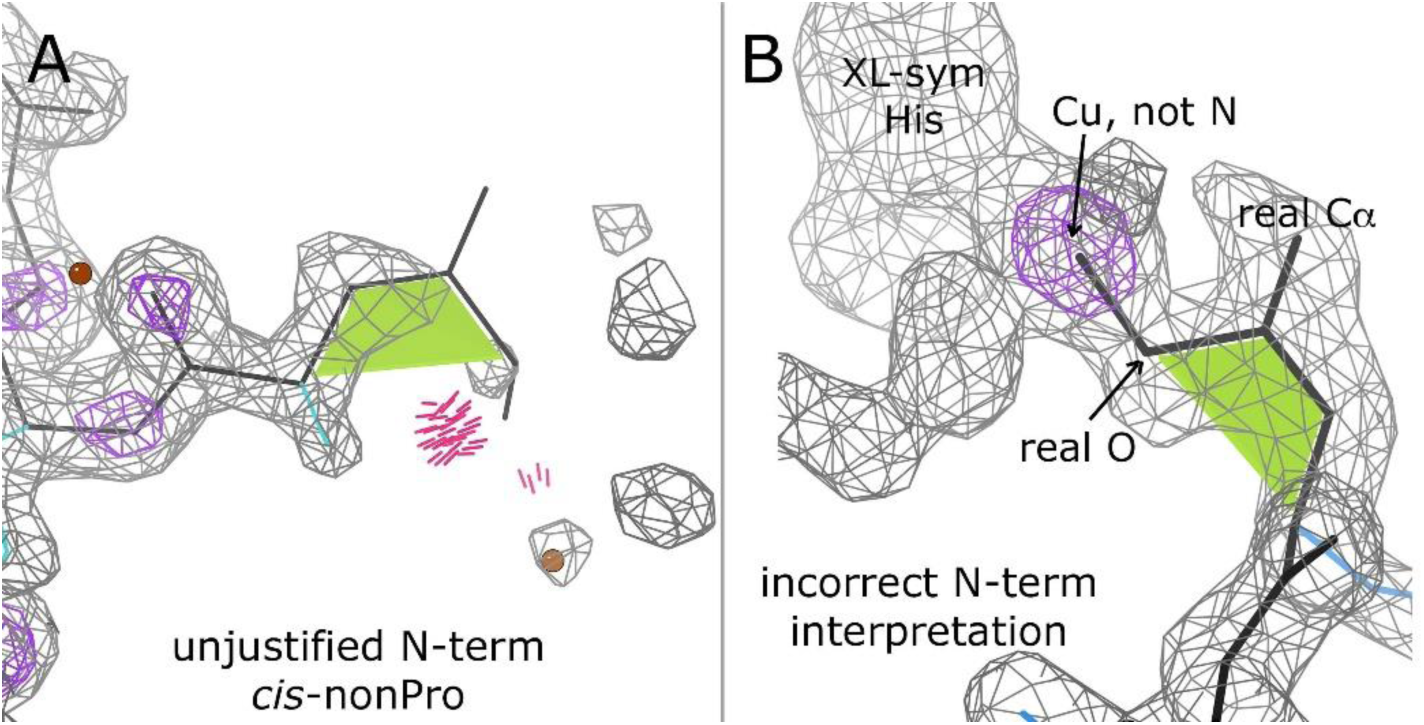
Non-proline *cis*peptides at chain temini (A) A typical misfit *cis*-nonPro at a chain N-terminus in 5Lp0. The truncated density encourages but does not justify fitting a *cis* conformation here. This residue is removed from the dataset for poor fit to map and a clash. (B) One of 3 incorrect N-terminal *cis*-nonPro that passed the filters. This is a case of the systematic-error switch between the carbonyl O and the Cα around the branch at the carbonyl C, but is very unusual in having strong density for the actual CO that is a ligand to an adventitious, crystal-contact Cu site in this copper-export protein (4f2f). Gray contour represents 1.2σ map level used by filters.

Residues with atoms that fail any of these criteria are removed from the PDB files entirely.

The fit-to-map criteria (B-factor, RSCC, and map value) are obtained using `phenix.real_space_correlation detail=atom`. This is done on the full parent structures before splitting into chains. Fit-to-map assessment could not be performed for 190 structures due to bad MTRIX records or other data issues. Chains from those structures were discarded. The B-factor, RSCC, and map cutoffs are those developed during production of a rotamer library using our previous, Top8000 dataset^2^.

Residues with alternate conformations are removed because the split in electron density that indicates an alternate necessarily lowers both those electron density peaks and the certainty of the model, especially when the alternates criss-cross each other, as frequently true. For applications that require the presence of alternates, we recommend that alternate conformations be validated and filtered as ensembles, taking care to spread alternate status along the backbone and to other interacting structure as needed to preserve acceptable geometry and avoid serious clashes.

Chains which were < 60% complete after residue filtering were discarded from the final dataset. This serves as a final check on overall structure quality and reduces the amount of chain fragmentation in the included chains. Mainchain-filtered chains typically retain a high level of completeness after filtering (Figure 1B), with almost half of the chains at least 90% complete. Full-residue-filtered chains are less complete, as expected, but over half of the chains are at least 75% complete after filtering.

At these high resolutions the protein core is well resolved and the disordered, poor-density stretches that fail the residue-level filtering are almost all at the surface and short. This means that the accepted continuous fragments between gaps are quite long and thus suitable as a basis of fragment libraries for model building, protein design or prediction, loop completion, and other uses. After filtering on mainchain atoms, 70% of remaining residues are in continuous segments of at least 15 residues, and 47% are in very long segments of at least 30 residues. Full-residue filtering is more sensitive and the results are more fragmented, with 31% of residues in segments of at least 15, and only 10% in segments of at least 30.

Only protein residues were filtered. Individual filtering of ligands, ions, and waters is beyond the current scope of this dataset. Ligands, ions, and waters are included in these files in the interest of completeness, but no guarantee of their quality is implied.

This residue-filtering filtering process sounds rather like the model validation that happens later, but the two procedures have distinct requirements. Reference-data filtering needs to be both stricter than, and independent of, the conformational model-validation criteria they will be used to define (such as rotamer, Ramachandran, or CaBLAM outliers). Local map quality is especially critical in reference data, because of the need to ensure that the experimental data is sufficient to specify the modeled conformation.

### In-file Documentation

The results of residue-level filtering are documented in each resulting PDB file by USER records appended to the end of the file. The necessity for including such documentation in the files themselves is why we use PDB format rather than the now-standard CIF format, which does not (yet) support USER__ records or an equivalent. These records report: 1) the residues that were removed and the reasons for their removal (as a string of 6 single-letter codes), 2) the residues that remain and the lengths of the sequence fragments they form, and 3) the overall completeness statistics for the filtered file. These USER records are self-documented, as in this small sample from 6git^12^:

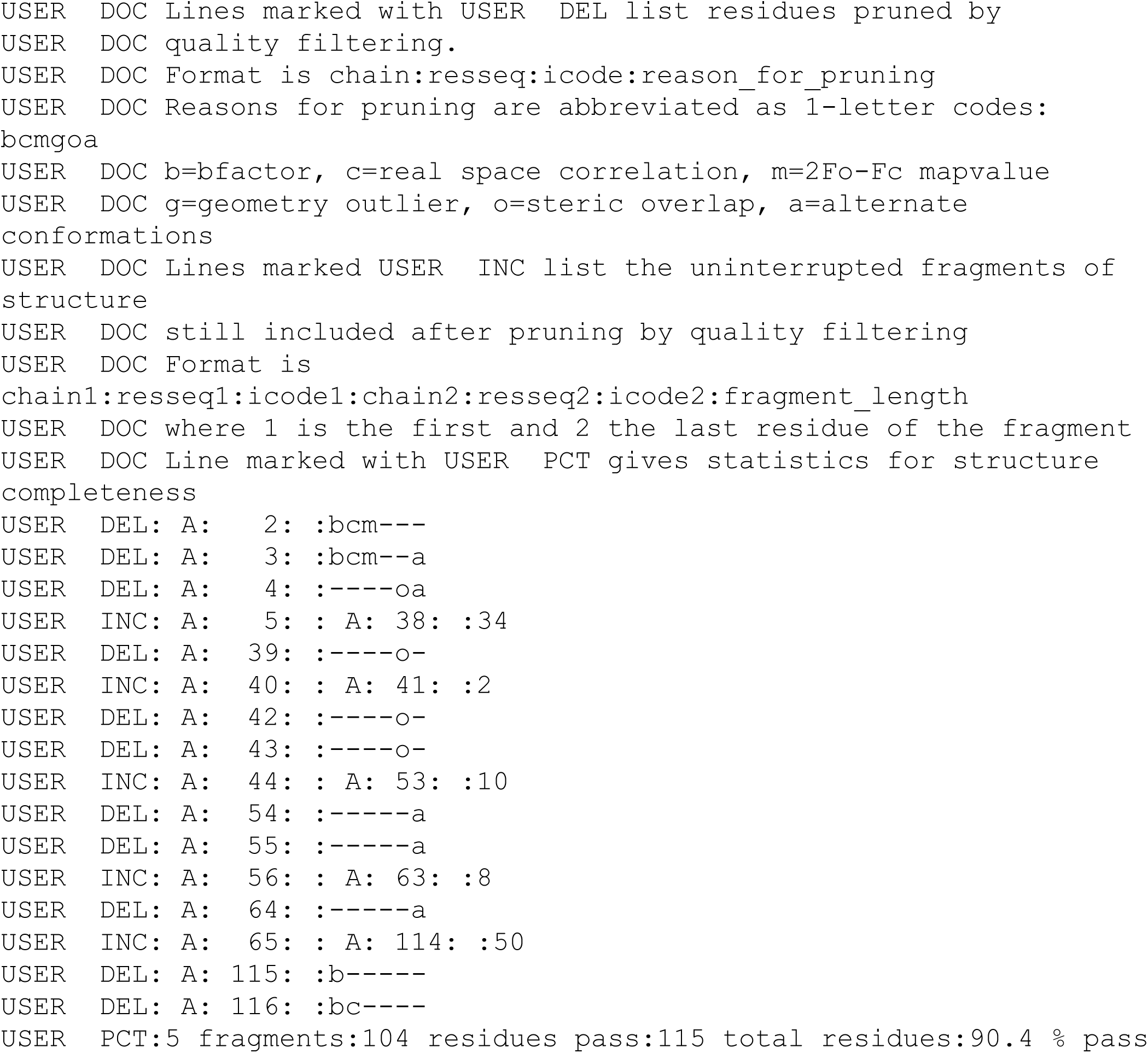

## Results and Causes of Filtering

Our previous dataset, the Top8000, contained 7957 chains at the 70% sequence identity level and just over 1 million residues after full-residue filtering. Table I shows that the Top2018 contains about as many chains and more residues even at its most stringent level (30% sequence identity). At the equivalent 70%, there are now 1.5 times as many chains and 3 times as many residues. This relative increase in accepted residues per chain indicates an increase in structure size and in general structure quality^13^ since 2013.

**Table I:** Chain and residue counts in the Top2018 datasets

|  | Mainchain-filtered chains | Mainchain-filtered residues | Full-residue-filtered chains | Full-residue-filtered residues |
| --- | --- | --- | --- | --- |
| 90% sequence<br>identity | 15,182 | 3,303,037 | 13,456 | 2,677,985 |
| 70% sequence<br>identity | 13,677 | 2,997,434 | 12,125 | 2,434,475 |
| 50% sequence<br>identity | 11,806 | 2,561,093 | 10,418 | 2,073,991 |
| 30% sequence<br>identity | 8,307 | 1,734,509 | 7,237 | 1,584,102 |

Table IIA shows the total contributions of each residue-level filtering criterion to the overall filtering. Bfactor is the dominant criterion, followed by alternate conformations, 2mFo-DFc map value, and clashes. Map value is more important for full-residue filtering due to the relative abundance of unresolved sidechains. Covalent geometry contributes the fewest total filterings, since this geometry is often tightly restrained in refinement. Nevertheless, geometry outliers >4σ usually diagnose severe modeling problems and are no less important for their rarity.

**Table IIA:** Percent of filtered-out residues that fail each criterion. A residue can fail multiple criteria.

|  | B factor | RSCC | map value | geometry | clash | alternates |
| --- | --- | --- | --- | --- | --- | --- |
| Mainchain<br>filter | 54% | 13% | 22% | 3% | 19% | 24% |
| Full-residue<br>filter | 52% | 24% | 55% | 3% | 28% | 14% |

Table IIB shows the unique contributions of each residue-level filtering criterion. B factor is again dominant, with 33% of filtered-out residues being removed due to high B factor and no other causes. Alternate conformations and clashes are the other largest independent contributors.

**Table IIB:**
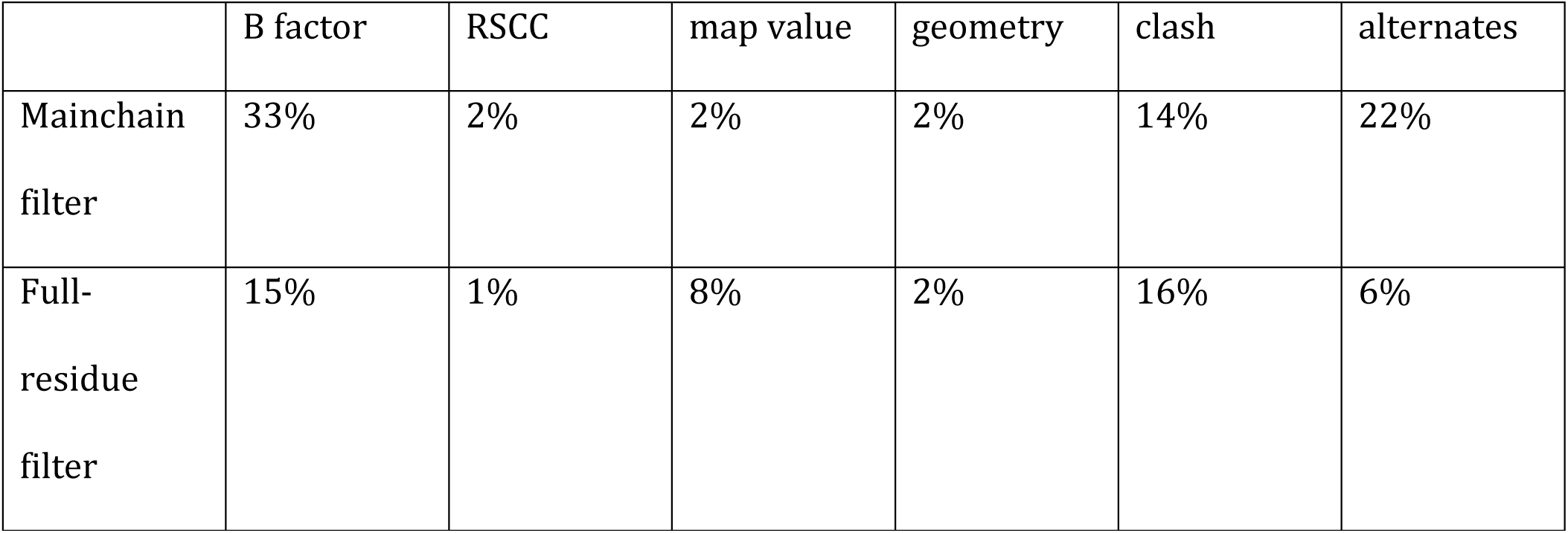
Percent of pruned residues uniquely pruned by each criterion

Residues which failed on real space correlation coefficient or map value criteria usually also failed on at least one other measure.

## Limitations of residue filtering

There is no substitute for visual inspection of structures. Our earliest dataset, the Top100, was hand curated. However, as the Protein Data Bank expanded, manual inspection of each structure became impractical. Automated quality assessments are required, but we still examine a sample of the results to ensure our automated code is doing what we intended.

Setting filtering cutoffs is a challenging balance. Overly-aggressive selection against outliers on datasets that are then used to define outliers is circular reasoning. Preserving real irregularities, where they are justified, is also important to understanding the true variety of protein structure. Our goal is to set cutoffs where they permit most of the justified conformations and eliminate most of the unjustified ones, with minimal false identifications leaking through in either direction. To achieve this, cutoffs are necessarily set in the ambiguous region between clearly good and clearly bad structure. Our identification of outliers for validation is set somewhat toward the forgiving side, to minimize “false alarms” for the structural biologist seeking to correct problems. The residue-level filtering cutoffs in this reference data are more toward the strict side, to minimize inclusion of incorrect local conformations. There is no magical, perfect cut within the ambiguous region.

An important commonality between reference-data filtering and later validation is that there are always at least a few genuine examples well-supported by their experimental data but outside the prior-probability expectations from previous structures. Those examples can be correct, appropriate for reference data, but are still outliers (e.g., non-rotameric sidechains). As well as having support in the map density, strained conformations need to be held in place. A couple of H-bonds can constrain one eclipsed dihedral for a polar sidechain, and tight packing interactions can constrain an otherwise-unfavorable conformation of a nonpolar sidechain, especially an aromatic ring^14^.

Our residue-level filtering relies heavily on fit-to-map criteria. This is appropriate for keeping our filtering independent of the conformation-based validations we use these datasets to construct. However, as a result, some types of systematic errors that are not strongly marked by poor map fit are not fully excluded from this dataset, regardless of filtering.

Leucine has a systematic-error conformation (Figure 2A) that can evade these filters often enough to produce a datapoint cluster (Figure 2B, pink dots); that cluster is an “imposter” rotamer that should not be included in a rotamer library. In this modeling error, χ2 is rotated by 140-150° and χ1 by 30-40° from the very common **tp** (trans, plus) conformation, which keeps the Cδ atoms in about the same peaks^14^. This is clearly an attractive but incorrect fit to the density, and both branches are almost exactly eclipsed. But the sidechain atoms sometimes fit within the density envelope just well enough to pass our filters. Fit-to-map filters aggressive enough to catch and remove all such tetrahedral-branch “imposters” would also remove real but marginal conformations that we wish to preserve.

## Importance of Residue Filtering

The key fact that motivates preparation of these datasets is that good average model quality across a whole structure is nevertheless compatible with extremely bad model quality in locally disordered regions with poor density. Familiar cases of this are mobile, unresolved sidechains on a protein’s surface compared to well-packed sidechains in a protein’s core, and unseen backbone at chain termini or in disordered loops. Chain-level filtering is an important first step, but it does not provide protection against residue-level modeling errors in those disordered regions of poor density.

The general-case Ramachandran distribution for residues in the Top2018 70% sequence identity dataset before residue-level filtering (Figure 3A) shows blurred edges and many outliers in the excluded regions. Filtering on mainchain atoms (Figure 3B) renders the edges of the distribution cleaner and more interpretable and removes many outliers. The difference in the distribution without filtering is sufficient to change the 0.2% contour boundary between allowed and outlier conformations (Figure 3C), by a greater amount than does the omission of all residues in secondary structures.

Sidechain rotamer distributions are even more affected. For example, in isoleucine, significant populations of **pm** and **tm** highly strained conformations (which clash with their own backbone) are present in the unfiltered set (Figure 3D). Full-residue filtering removes only 15-25% of the 5 most common Ile rotamers, while it removes 70-80% of these 2 disfavored conformations (Figure 3E), placing their remaining populations below the statistical threshold for being identified as true rotamers (Figure 3F).

We examined all 13 examples of Ile **tm** that passed the filters, finding 12 of them clearly supported by map density and constrained into this unfavorable conformation by tight packing that cannot accommodate any other rotamer. In one case (5nz4^15^ Ile B 216) the packing allows a non-clashing **tp** rotamer, but it is not represented in the weak density even at a very low contour level. This check confirms that even for extremely rare conformations, examples in the filtered data are likely to be correct. Figure 4A shows one of these genuine conformations with completely unambiguous density peaks for each atom and very good all-atom contacts (Ile 103 in 4ezi^16^). The local necessity for this rare and strained conformation is explained by Figure 4B, which shows in orange the geometrically “ideal” **tm** conformation at -180°, -60° that would have a very bad clash of Cδ and its hydrogens with the following peptide. To avoid that clash, χ2 has to change by 28° and each of the 3 covalent angles from the backbone out to Cδ is opened up by about 2°, a large total amount of strain. No more favorable conformation is possible for this sidechain because it is closely surrounded by other structure, especially the tyrosine packed tightly above it.

As another telling example of misfitting a rare, strained conformation, the structural biology community may remember the crisis of gross *cis*-non-proline over-use from about 2006 until after the problem was recognized^17,18^. This phenomenon was most extreme at lower resolutions (often >1% of total residues, or 30 times too many), but is also present in poorly-resolved regions of high-resolution structures such as those in this dataset. The over-use was largely cured just by obvious flagging of *cis*-nonPro as probable outliers in graphics or validation reports^19,20^. Residue-level filtering of our reference data has always guarded against the inclusion of unsupported *cis*-nonPro peptides, on both a statistical and an individual level.

Before filtering, the 70% homology set of the Top2018 contained 1959 *cis*-nonPro out of 3,324,246 evaluable peptide bonds, for an occurrence rate of 0.048% or about 1 in 2000 (a rate often reported before any data-quality controls). After filtering, there remain 776 *cis*-nonPro out of 2,652,118, for an occurrence rate of 0.029% or about 1 in 3500. This lower rate agrees with recent analyses of valid *cis*-nonPro occurrence (see accompanying paper).

More importantly than these general statistics, residue-level filtering removes nearly all of the obviously incorrect *cis*-nonPro peptides from the dataset. These include some known, systematic patterns of incorrect *cis* modeling, such as building *cis*-peptides into the weak, patchy, or truncated density at chain termini or in partly disordered loops (Figure 5A, 4rm4^21^ 170-172). The lack of strong electron density in such regions *allows* this and other modeling errors to occur. Thus it is vital to the health of a statistical reference dataset or fragment library to remove these regions of low certainty, as we do in this dataset. In contrast, real *cis*-nonPro are supported by clear electron density (Figure 5B, 6btf^22^)

As a specific case, we considered *cis*-nonPro peptides at the very vulnerable position of chain ends (both the first and last residues in the chain, and the residues at the ends of unmodeled loops). The unfiltered Top2018 data contains 200 chain-terminal instances of *cis*-nonPro. Most of these 200 cases occur at the N-rather than C-terminus of a modelled stretch, where the lack of preceding structure makes the choice vulnerable to a misfit *cis* rather than *trans* peptide at the branch point between the carbonyl O and the N-terminal Cα (Figure 6A, 5Lp0). 197 (98.5%) of them fail the filters and are removed. Of the 3 (1.5%) that passed the filters, all three are made difficult to avoid or validate by an N-terminal Gly that provides no sidechain to help choose the correct alternative, and two are in rather weak, ambiguous density that only barely passes the filters. The third example is actually an adventitious crystal-contact Cu site (4f2f^23^ Gly-Ser 1), where the Cu is incorrectly modeled as the N-terminal N (Figure 6B). The density peak is >16σ and the B-factor of the modeled N is just 5 compared to B-factors of 20-30 for the other atoms of Gly 0, both indications that a much heavier atom should have been modeled there.

Residue-level filtering thus ensures that the population of *cis*-nonPro peptides is not statistically or locally overrepresented due to modelling errors. The *cis*-nonPro that remain in the dataset (Figure 5B) do so based on a reasonable standard of map and model quality and provide genuine *cis* conformations and contexts that can be used in model building when the evidence in the experimental data is good enough to outweigh the very low prior probability of 1:3500 (8 log likelihood units). If alternative *trans* and *cis* conformations are explicitly compared, the choice is usually clear at better than about 2.3Å. At resolution ≥ 3Å or when the density is otherwise ambiguous, the much less probable *cis* alternative should never be chosen, unless it occurs in a related protein at higher resolution (see accompanying paper for more details).

## Conclusions

Residue-level filtering is necessary even in otherwise excellent structures, in order to remove significant populations of disordered residues whose conformations have little or no experimental support. Conclusions based on unfiltered residues will be inaccurate representations of true protein behavior in both their statistics and their details. The easy availability of this pre-filtered reference dataset on Zenodo is therefore an important step forward for structural bioinformatics.

The full-coordinate, residue-filtered reference datasets described here omit all residues that fail the quality filters, so that they contain only coordinates for residues which are almost certainly correct. These gapped, residue-filtered datasets are suitable for most uses, though not for applications that require the full, ungapped context, such as Voronoi analyses or molecular dynamics simulations. The full-residue quality-filtered reference dataset can be used to prepare protein sidechain rotamer libraries^2,14^ or to study macromolecular structural motifs that span multiple residues and involve backbone-sidechain interactions^24,25^. The mainchain residue-filtered reference dataset can be used to define Ramachandran distributions^8,26^, to study rare backbone conformations such as *cis*-nonPro^20^, and to prepare curated fragment libraries for model-building or for protein design^13,18,27^.

Future directions for this work include compiling a similar reference dataset for RNA structures, developing advanced map-quality criteria to consider atom type and use both upper and lower cutoffs, and responding to user feedback for what expansions would help enable other unanticipated uses.

## Availability

These datasets are available on the Zenodo data repository, each at four levels of sequence redundancy.

The mainchain-filtered set is here: https://doi.org/10.5281/zenodo.4626149.

The full-residue-filtered set is here: https://doi.org/10.5281/zenodo.5115232.

Zenodo supports versioning, and these links will resolve to the latest version of each dataset.

## Acknowledgements

This work was funded by NIH grants R35-GM131883 to DCR and P01-GM063210 Project IV to JSR. The authors declare no conflicts of interest.

